# Exploration in the wild

**DOI:** 10.1101/492058

**Authors:** Eric Schulz, Rahul Bhui, Bradley C. Love, Bastien Brier, Michael T. Todd, Samuel J. Gershman

## Abstract

Making good decisions requires people to appropriately explore their available options and generalize what they have learned. While computational models have successfully explained exploratory behavior in constrained laboratory tasks, it is unclear to what extent these models generalize to complex real world choice problems. We investigate the factors guiding exploratory behavior in a data set consisting of 195,333 customers placing 1,613,967 orders from a large online food delivery service. We find important hallmarks of adaptive exploration and generalization, which we analyze using computational models. We find evidence for several theoretical predictions: (1) customers engage in uncertainty-directed exploration, (2) they adjust their level of exploration to the average restaurant quality in a city, and (3) they use feature-based generalization to guide exploration towards promising restaurants. Our results provide new evidence that people use sophisticated strategies to explore complex, real-world environments.

## Introduction

When facing a vast array of new opportunities, a decision maker has two key tasks: to acquire information (often through direct experience) about available options, and to apply that information to assess options not yet experienced. These twin problems of *exploration* and *generalization* must be tackled by any organism trying to make good decisions, but they are challenging to solve because optimal solutions are computationally intractable^1^. Consequently, the means by which humans succeed in doing so—especially in the complicated world at large—have proven puzzling to psychologists and neuroscientists. Many heuristic solutions have been proposed to reflect exploratory behavior^2–4^, inspired by research in machine learning^5^. However, most empirical studies have used a small number of options and simple attributes^6^. To truly ascertain the limits of exploration and generalization requires empirical analysis of behavior in the world outside the lab.

We study learning and behavior in a complex environment using a large data set of human foraging in the “wild”—online food delivery. Each customer has to decide which restaurant to pick out of hundreds of possibilities. How do they make a selection from this universe of options? Guided by algorithmic perspectives on learning, we look for signatures of adaptive exploration and generalization that have been previously identified in the lab. This allows us not only to characterize these phenomena in a naturally incentivized setting with abundant and multi-faceted stimuli, but also to weigh in on existing debates by testing competing theories of exploratory choice.

We address two broad questions. First, how do people strategically explore new options of uncertain value? Different algorithms have been proposed to describe exactly how uncertainty can guide exploration in qualitatively different ways, such as by injecting *randomness* into choice, or by making choices *directed* toward uncertainty^2, 3, 7^. However, results have been mixed, and these phenomena remain to be studied under real-world conditions. Second, how do people generalize their experiences to other options? Modern computational theories make quantitative predictions about how feature-based similarity should govern generalization, which can in turn guide choice. But again it is unclear whether these theories can successfully predict real-world choices.

Our results indicate that customers explore (i.e., order from unexperienced restaurants) adaptively based on signals of restaurant quality, and make better choices over time. Exploration is indeed risky and leads to worse outcomes on average, but people are more willing to explore in cities where this downside is lower due to higher mean restaurant quality. Importantly, we show that customers’ exploratory behavior takes into account not only the prospective reward from choosing a restaurant, but also the degree of uncertainty in their reward estimates. Consistent with an optimistic uncertainty-directed exploration policy, they preferentially sample lesser known options and exhibit higher sampling entropy after a bad (compared to a good) outcome.

We find that choices are best fit by a model that includes both an “uncertainty bonus” for unfamiliar restaurants, and a mechanism for generalization by function learning (based on restaurant features). People appear to benefit from such generalization, as exploration yields better realized outcomes in cities where features have more predictive power. We also show that people generalize their experiences across different restaurants within the same broad cuisine type, defined both empirically within the data set, and by independent similarity ratings. As predicted by a combination of similarity-based generalization and uncertainty-directed exploration, good experiences encourage selection of other restaurants within the same category, while bad experiences discourage this to an even greater extent.

In order to set the stage for our analyses of purchasing decisions, we first review the algorithmic ideas that have been developed to explain exploration in the laboratory.

## Prior work on the exploration-exploitation dilemma

### Uncertainty-guided algorithms

Most of what we know about human exploration comes from *multi-armed bandit tasks*, in which an agent repeatedly chooses between several options and receives reward feedback^8, 9^. Since the distribution of rewards for each option is unknown at the beginning of the task, an agent is faced with an *exploration-exploitation dilemma* between two types of actions: should she exploit the options she currently knows will produce high rewards while possibly ignoring even better options? Or should she explore lesser-known options to gain more knowledge but possibly forego high immediate rewards? Optimal solutions only exist for simple versions of this problem^1^. These solutions are in practice difficult to compute even for moderately large problems. Various heuristic solutions have been proposed. Generally, these heuristics coalesce around two algorithmic ideas^10, 11^. The first one is that exploration happens randomly, for example by occasionally sampling one of the options not considered to be the best^12^; or by so-called soft-maximization of the expected utilities for each option—i.e., randomly sampling each option proportionally to its value. The other idea is that exploration happens in a directed fashion, whereby an agent is explicitly biased to sample more uncertain options. This uncertainty-guidance is frequently formalized as an “uncertainty bonus”^5, 13^ which inflates an option’s expected reward by its uncertainty.

There has been a considerable debate about whether or not directed exploration is required to explain human behavior^14, 15^. For example, Daw and colleagues^14^ have shown that a softmax strategy explains participants’ choices best in a simple multiarmed bandit task. However, several studies have produced evidence for a direct exploration bonus^4, 16, 17^. Recent studies have proposed that people engage in both random and directed exploration^2, 7, 18, 19^. It has also been argued that directed exploration might play a prominent role in more structured decision problems^20, 21^. However, evidence for such algorithms is still missing in real-world purchasing decisions^6^.

### Generalization

Multiple studies have emphasized the importance of generalization in exploratory choice. People are known to leverage latent structures such as hierarchical rules^22^ or similarities between a bandit’s arms^23^.

Gershman et. al^24^ investigated how generalization affects the exploration of novel options using a task in which the rewards for multiple options were drawn from a common distribution. Sometimes this common distribution was “poor” (options tended to be non-rewarding), whereas sometimes the common distribution was “rich” (options tended to be rewarding). Participants sampled novel options more frequently in rich environments than in poor environments, consistent with a form of adaptive generalization across options.

Schulz et al.^25^ investigated how contextual information (an option’s features) can aid generalization and exploration in tasks where the context is linked to an option’s quality by an underlying function. Participants used a combination of functional generalization and directed exploration to learn the underlying mapping from context to reward (see also^21, 26^).

## Results

We looked for signatures of uncertainty-guided exploration and generalization in a data set of purchasing decision taken from the online food delivery service *Deliveroo* (see Materials and Methods for more details). Further analyses and method details can be found in the Supplemental Information. In the first two sections of the Results, we provide some descriptive characterizations of the data set. In particular, we show that customers learn from past experience and adapt their exploration behavior over time. Moreover, their exploration behavior is systematically influenced by restaurant features and hence amenable to quantification. We then turn to tests of our model-based hypotheses. We find that customers’ exploration behavior can be clustered meaningfully, exhibits several signatures of intelligent exploration which have previously been studied in the lab, and can be captured by a model that generalizes over restaurant features while simultaneously engaging in directed exploration.

### Learning and exploration over time

We first assessed if customers learned from past experiences, as reflected in their order ratings over time (Fig. 1a). The order rating is defined as customers’ evaluation on a scale between 1 (poor) and 5 (great). Customers picked better restaurants over time: there was a positive correlation between the number of a customer’s past orders and her ratings (*r* = 0.073; 99.9% CI: 0.070, 0.076).

**Figure 1.**
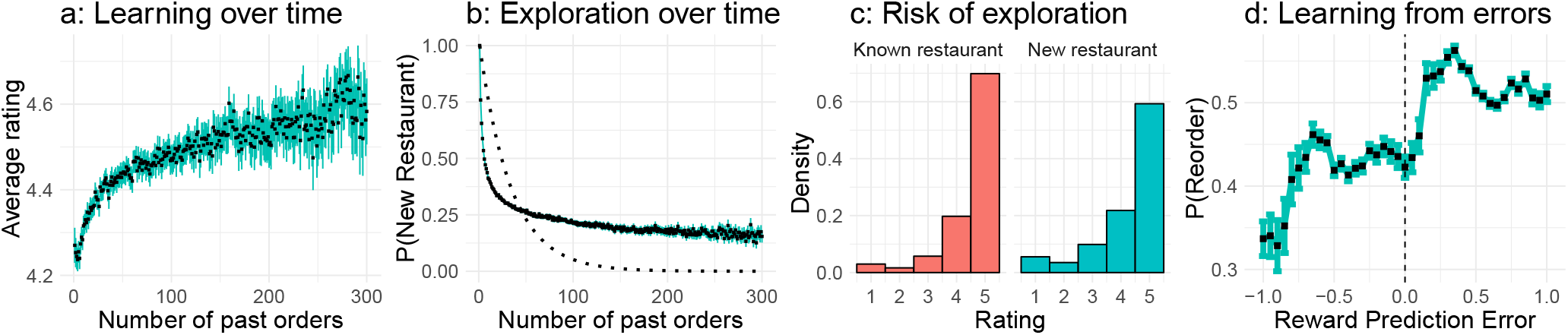
Learning and exploration over time. **a:** Average order rating by number of past orders. **b:** Probability of sampling a new restaurant in dependency of the number of past orders. Dashed black line indicates simulated exploration behavior of agents randomly exploring available restaurants. **c:** Distribution of order ratings for newly sampled and known restaurants. **d:** Average probability of reordering from a restaurant as a function of reward prediction error. Means are displayed as black squares and error bars show the 95% confidence interval of the mean.

Next, we assessed exploratory behavior by creating a variable indicating whether a given order was the first time a customer had ordered from that particular restaurant—i.e., a signature of pure exploration^24, 26^. Figure 1b shows the averaged probability of sampling a new restaurant over time (how many orders a customer had placed previously).

Customers sampled fewer new restaurants over time, leading to a negative overall correlation between the number of past orders and the probability of sampling a new restaurant (*r* = −0.139; 99.9% CI: −0.142, −0.136). Exploration also comes at a cost (Fig. 1c), such that explored restaurants showed a lower average rating (mean rating=4.257, 99.9% CI: 4.250, 4.265) than known restaurants (mean rating=4.518, 99.9% CI: 4.514, 4.522).

Customers learned from the outcomes of past orders. Figure 1d shows their probability of reordering from a restaurant as a function of their reward prediction error (RPE; the difference between the expected quality of a restaurant, as measured by the restaurant’s average rating at the time of the order, and the actual pleasure customers perceived after they had consumed the order, as indicated by their own rating of the order). RPEs are a key component of theories of reinforcement learning^27^, and we therefore expected that customers would update their sampling behavior after receiving either a positive or a negative RPE. Confirming this hypothesis, customers were more likely to reorder from a restaurant after an experience that was better than expected (positive RPE: p(reorder)=0.518, 99.9%; CI: 0.515, 0.520) than after an experience that was worse than expected (negative RPE: p(reorder)=0.394, 99.9%; CI: 0.391, 0.398). The average correlation between RPEs and the probability of reordering was *r* = 0.110 (99.9% CI: 0.107, 0.114).

### Determinants of exploration

In the next part of our analysis, we focused on what factors influenced customers’ decisions to explore a new restaurant. In particular, we assessed if exploration behavior was systematic and therefore looked at the following four restaurant features that were always visible to customers at the time of their order: the relative average price of a restaurant, its standardized estimated delivery time, the mean rating of a restaurant at the time of the order, and the number of people who had rated the restaurant before.

Customers preferred restaurants that were comparatively cheaper (Fig. 2a): the correlation between relative price and the probability of exploration was negative (*r* = −0.059; 99.9% CI: −0.0641, −0.0548). There was a non-linear relationship between a restaurant’s estimated delivery time and its probability of being explored (Fig. 2b): exploration was most likely for standardized delivery times between 1 and 2.5 (0.288, 99.9% CI: 0.285, 0.292), and less likely for delivery times below 1 (0.288, 99.9% CI: 0.285, 0.292 or above 2.5 (0.252, 99.9% CI: 0.229, 0.274). This indicates that customers might have taken into account how long it would take to plausibly prepare and deliver a good meal when deciding which restaurants to explore. The average rating of a restaurant also affected customers’ exploratory behavior (Fig. 2c): higher ratings were associated with a higher chance of exploration (*r* = 0.038; 99.9% CI: 0.0337, 0.0430). The number of ratings per restaurant also influenced exploration (Fig. 2d), with a negative correlation of *r* = −0.188 (99.9% CI: −0.192, −0.183). This correlation is relatively large because restaurants that have been tried more frequently are less likely to be explored for the first time. We therefore repeated this analysis for all restaurants that had been rated more than 500 times, yielding a correlation of *r* = −0.034 (99.9% CI: −0.042, −0.026).

**Figure 2.**
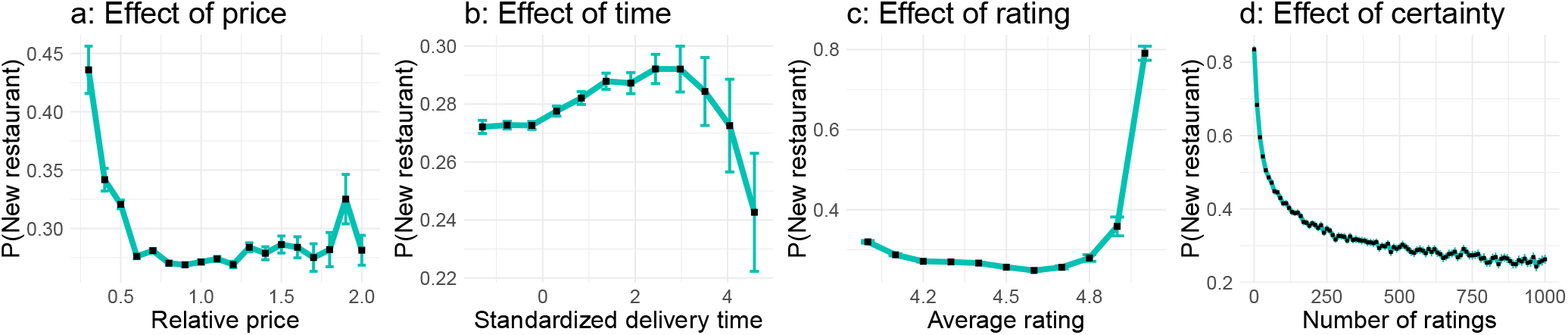
Factors influencing exploration. **a:** Effect of relative price. The relative price indicates how much cheaper or more expensive a restaurant was compared to a median restaurant in the same city. **b:** Effect of standardized (z-transformed) estimated delivery time. **c:** Effect of average rating. **d:** Effect of a restaurant’s number of past ratings (certainty). Means are displayed as black squares and error bars show the 95% confidence interval of the mean.

**Table 1.** Results of the mixed-effects logistic regression.

|  | Estimate | Std. Error | z value | Pr(>\|z\|) |
| --- | --- | --- | --- | --- |
| Intercept | -0.663 | 0.008 | -82.01 | <.001 |
| Relative price | -0.014 | 0.006 | -2.27 | .02 |
| Time-Linear | -0.0246 | 0.008 | -3.22 | .001 |
| Time-Quadratic | 0.015 | 0.004 | 3.89 | <.001 |
| Average rating | 0.086 | 0.006 | 13.85 | <.001 |
| Number of ratings | -0.475 | 0.007 | -70.27 | <.001 |

We standardized and entered all of the variables into a mixed-effects logistic regression modeling the exploration variable as the dependent variable and adding a random intercept for each customer (see SI for full model comparison). We again found that a smaller number of total ratings (*β* = −0.475), a higher average rating (*β* = 0.086), and a lower price (*β* = −0.014) as well as a quadratic effect of time (*β*_Linear_ = −0.025, *β*_Quadratic_ = 0.015) were all predictive of customers’ exploration behavior. In summary, exploration in the domain of online ordering is systematic, interpretable and amenable to quantification. We next turned to an examination of our model-based hypotheses concerning uncertainty-directed exploration and generalization.

## Signatures of uncertainty-directed exploration

We probed the data for signatures of uncertainty-directed exploration algorithms that attach an uncertainty bonus to each option. One such signature is that directed and random exploration make diverging predictions about behavioral changes after either a positive or a negative outcome. Whereas random (softmax) exploration predicts no difference in sampling behavior after either a better or a worse-than-expected outcome, directed exploration predicts a stronger increase in sampling behavior after a worse-than-expected outcome (see SI). This is due to the properties of algorithms that assess an option’s utility by a weighted sum of its expected reward and its standard deviation. After a bad experience, the mean and standard deviation both go down, whereas after a good experience the mean goes up but the standard deviation goes down. Thus, there should be more changes in customers’ sampling behavior after a bad than after a good outcome.

We verified this prediction by calculating the Shannon entropy of customers’ next 4 purchases after having experienced either a better-than or a worse-than-expected order. The calculated entropy was higher for negative RPEs (Fig 3a; 1.112, 99.9% CI: 1.109, 1.115) than for positive RPEs (1.082, 99.9% CI: 1.081, 1.084), in line with theoretical predictions of a directed exploration algorithm. This difference was present throughout time, such that the effect size when comparing the entropies for negative and positive RPEs for different numbers of past orders (of a given customer) always revealed a negative effect (Fig 3b). However, this effect decreased over time with a negative correlation between the effect and customers’ past orders of *r* = −0.68 (99.9% CI: −0.90, -0.33). This means that customers learned over time, leading to lower entropies in their sampling behavior after unexpected outcomes as they gained more experience.

**Figure 3.**
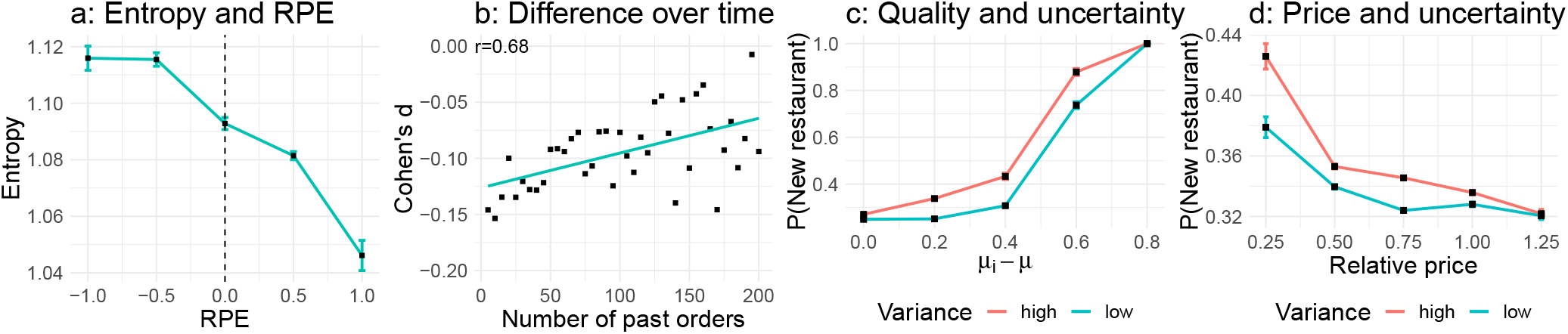
Signatures of uncertainty-directed exploration. **a:** Entropy of the next 4 choices in dependency of reward prediction error (RPE). **b:** Differences (signed Cohen’s d) in entropy for negative vs. positive RPEs over learning history. Turquoise line marks least-square regression line. **c:** Probability of choosing a novel restaurant in dependency of its difference to an average restaurant within the same cuisine type for restaurants with high and low relative variance. **d:** Probability of choosing a novel restaurant in dependency of its relative price for restaurants with high and low relative variance.

We assessed customers’ exploration behavior in dependency of the differences in ratings for a given restaurant as compared to the average of all restaurants within the same cuisine type (value difference). We also calculated each restaurant’s relative variance, i.e. how much more variance in its ratings a restaurant possessed as compared to the average variance per restaurant within the same cuisine type. The probability of exploring a new restaurant increased as a function of the restaurant’s value difference (Fig. 3c; *r* = 0.05, 99.9% CI: 0.045, 0.056). Additionally, a restaurant’s relative variance also correlated with its probability of being explored (Fig. 3c; *r =* 0.05; 99.9% CI: 0.045, 0.056). Comparing restaurants with a high vs. low relative variance in their ratings (based on a median split) revealed a shift of the choice function towards the left. In other words, restaurants with higher relative uncertainty (0.344; 99.9% CI: 0.341, 0.349) are preferred to restaurants with lower relative uncertainty (0.319; 99.9% CI: 0.317, 0.321), as predicted by uncertainty-directed exploration strategies^2, 18^. This difference can also be observed when repeating the same analysis using a restaurant’s price (Fig. 3d): as restaurants get more expensive, they are less likely to be explored (*r* = −0.017; 99.9%CI: −0.023, −0.013). This function is again shifted for restaurants with higher relative uncertainty: given a similar price range, relatively more uncertain restaurants are more likely to be explored than less uncertain restaurants.

To further validate these findings, we fit a mixed-effects logistic regression, using the exploration variable as the dependent variable. For the independent variables, we used the mean difference in ratings between the restaurant and the average restaurant within the same cuisine type, a restaurant’s relative price, and its relative uncertainty (see Tab. 2). The average value difference (*β* = 0.114), the relative price *β* = −0.0876) and the relative uncertainty (*β* = 0.084) all affected a restaurants’ probability to be explored. Thus, even when taking into account a restaurant’s price and its ratings, customers still preferred more uncertain options. This provides strong evidence for a directed exploration strategy.

**Table 2.** Results of mixed-effects logistic regression.

|  | Estimate | Std. Error | z value | Pr(>\|z\|) |
| --- | --- | --- | --- | --- |
| Intercept | -0.342 | 0.007 | 45.81 | <.001 |
| Value difference | 0.114 | 0.0135 | 8.47 | <.001 |
| Relative price | -0.087 | 0.007 | -11.67 | <.001 |
| Variance difference | 0.084 | 0.003 | 24.13 | <.001 |

### Signatures of generalization

We assessed if customers employed generalization to guide their purchasing decisions. We first looked at patterns of consecutive explorations (how one exploratory choice predicted the next one). Specifically, we looked at 20 frequent cuisine types and assessed how much exploring a restaurant from one type predicted exploring a restaurant from another type using a simple regression, and repeating this analysis for all combinations of types. This analysis revealed clusters of cuisine types within customers’ exploratory behavior (see Fig. 4a).

**Figure 4.**
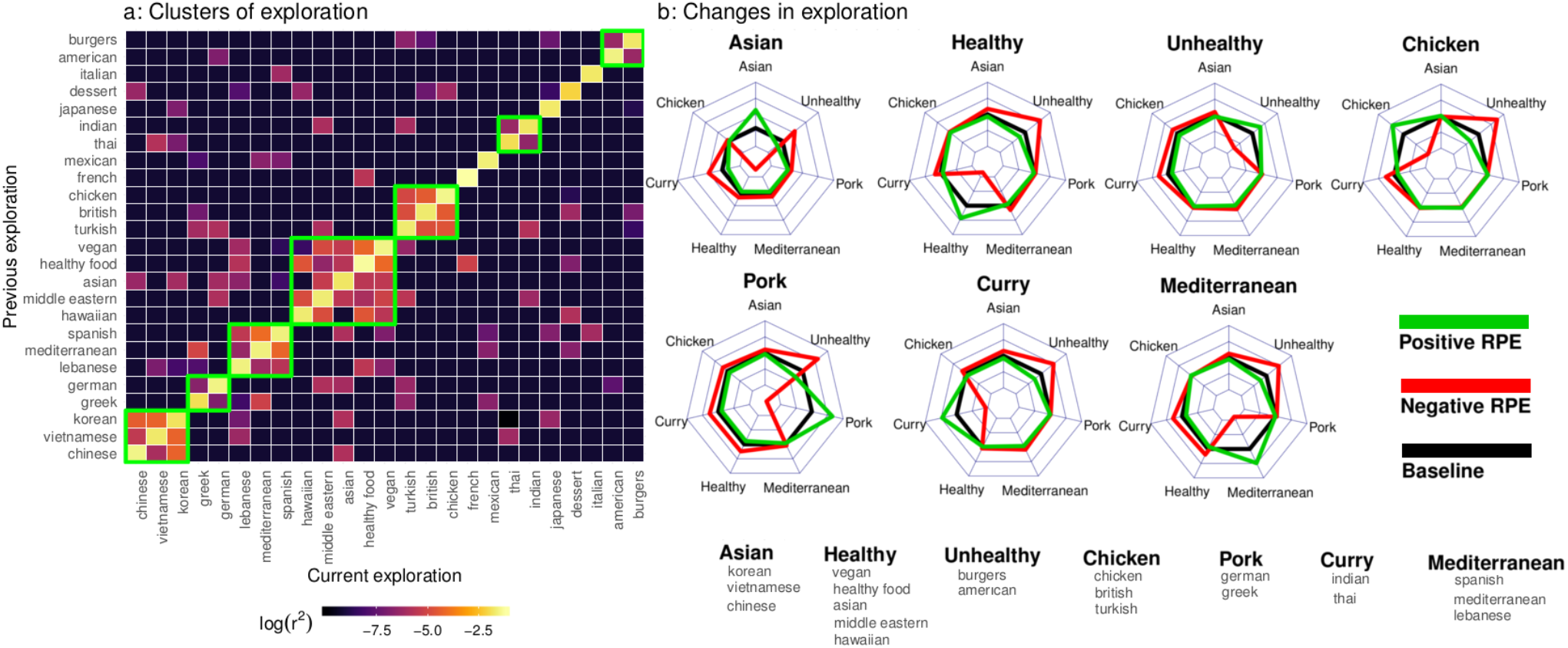
Signatures of direct exploration. **a:** Clusters of exploration between different cuisine types within customers’ consecutive explorations. Green rectangles mark clusters of exploration. **b:** Moves between clusters after better-than-expected (positive RPE) and worse-than-expected (negative RPE) outcomes as compared to a mean baseline. Centers of radar plots indicate a change of −5%, outermost lines indicate a change of +5%.

For example, exploring a restaurant from the cuisine type “Burgers” was predictive of exploring a restaurant from the cuisine type “American”, most of the Asian cuisine types clustered together, and so forth. As there were seven main clusters in total (see Materials and Methods and SI for details), this also allowed us to assess moves between different clusters after either a worse or a better-than-expected outcome. This led to four insights. First, customers were more likely to explore within the same cluster after a good outcome (+2.27%) and less likely after a bad outcome (−5.19%), while bad outcomes had a larger effect than good outcomes, again hinting at strategies of directed exploration. Secondly, customers’ switches between clusters were meaningful. For example, after a bad experience with the cluster “Asian”, customers frequently switched to the cluster “Curry” (+1.51%), which contained Indian and Thai cuisine. Thirdly, there were spill-over effects; for example, a positive experience with “Mediterranean” food also made exploring the cluster “Chicken” more likely (+0.5%). Finally, customers most often switched to exploring “Unhealthy” cuisine types after bad outcomes (+2.72%).

We also assessed if customers’ switches between different cuisine types could be related to a subjective understanding of similarity between types (Fig. 5a). We therefore asked 200 participants on Amazon’s Mechanical Turk to rate the similarity between 30 pairs of cuisine types sampled from the 20 above types on a scale from 0 (not at all similar) to 10 (totally similar). We then tested how much exploratory switches between cuisine types mapped onto the mean similarity ratings between cuisine types. There was a positive correlation between similarity ratings and the frequency of switches between cuisine types of *r* = 0.78. Thus, exploratory choices not only clustered into interpretable clusters, but were also predicted by subjective similarities between cuisine type.

**Figure 5.**
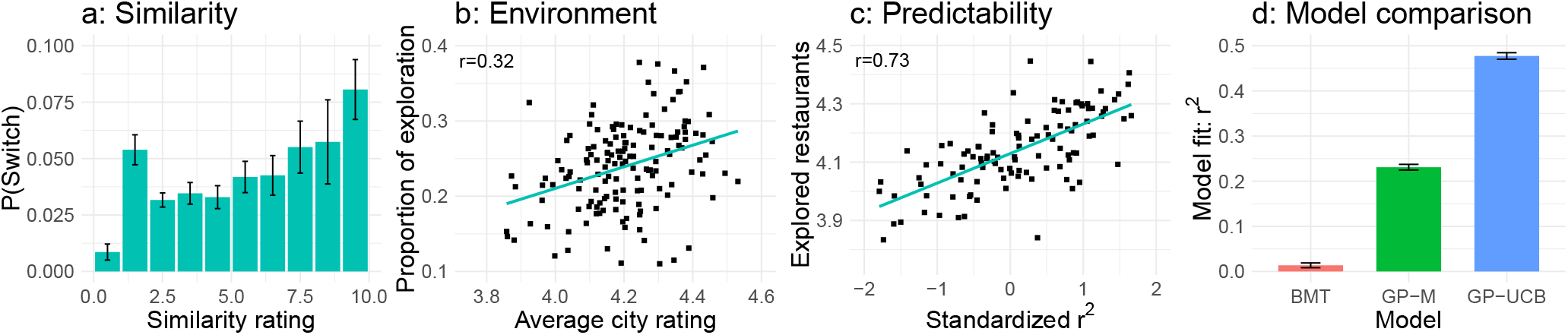
Signatures of generalization. **a:** Probability of switches between cuisine types and rated similarities between the same types. **b:** Average rating per city and proportion of exploratory choices. Turquoise line marks least-square regression line. **c:** Predictability of a restaurant’s quality and average rating of explored restaurants. Turquoise line marks least-square regression line. **d:** Results of model comparison for new customers’ behavior. Considered models were the Bayesian Mean Tracker (BMT), a Gaussian Process with a mean-greedy sampling strategy (GP-M), and a Gaussian Process with a Upper Confidence Bound sampling strategy (GP-UCB).

We further tested how much customers’ exploration was guided by generalization. Gershman et al.^24^ showed that participants explore novel options more frequently in environments where all options are generally good. We found evidence for this phenomenon in our data (Fig. 5b): there was a positive correlation between a city’s average restaurant rating and the proportion of exploratory choices in that city (*r* = 0.32; 99.9% CI: 0.21, 0.49, see SI for partial correlations).

Next, we examined whether customers’ explorations were more successful in cities where ratings were more predictable as assessed by how well individual ratings were predictable by the features’ price, delivery time, mean rating, and number of ratings using randomly sampled learning and tests sets of the same size for each city. Customers were more successful (i.e., gave higher ratings to sampled restaurants) in their exploratory choices in cities where ratings were generally more predictable (*r* = 0.73; Fig. 5d, 99.9% CI: 0.53, 0.84). Thus, customers took contextual features into account to guide their exploration, similar to findings in contextual bandit tasks^25, 26^.

In the attempt to test algorithms of both directed exploration and generalization simultaneously, we compared three models of learning and decision making based on how well they captured the sequential choices of 3,772 new customers who had just started ordering food and who had rated all of their orders. The first model was a Bayesian Mean Tracker (BMT) that does not generalize across restaurants, only learning about a restaurant’s quality by sampling it. The second model used Gaussian Process regression to learn about a restaurant’s quality based on the four observable features (price, mean rating, delivery time, and number of past ratings). Gaussian Process regression is a powerful model of generalization and has been applied to model how participants learn latent functions to guide their exploration^20, 21, 25^. This model was either paired with a mean-greedy sampling strategy (GP-M) or with a directed exploration strategy that sampled based on an option’s upper confidence bound (GP-UCB). We treated customers’ choices as the arms of a bandit and their order ratings as their utility, and then evaluated each model’s performance based on its one-step-ahead prediction error, standardizing performance by comparing to a random baseline. Since it was not possible to observe all restaurants a customer might have considered at the time of an order, we compared the different models based on how much higher in utility they predicted a customer’s final choice compared to an option with average features. The BMT model barely performed above chance (*r*^2^ = 0.013; 99.9% CI: 0.005, 0.022). Although the GP-M model performed better than the BMT model (*r*^2^ = 0.231; 99.9% CI: 0.220, 0.241), the GP-UCB model achieved by far the best performance (*r*^2^ = 0.477; 99.9% CI: 0.465, 0.477). Thus, a sufficiently predictive model of customers’ choices required both a mechanism of generalization (learning how features map onto rewards), and a directed exploration strategy (combining an restaurant’s mean and uncertainty to estimate its decision value).

## Discussion

We investigated customers’ exploration behavior in a large data set of online food delivery purchases. Customers learned from past experiences, and their exploratory behavior was affected by a restaurant’s price, average rating, number of ratings and estimated delivery time. Our results further provide strong evidence for several theoretical predictions: people engaged in uncertainty-directed exploration, and their exploration was guided by similarity-based generalization. Computational modeling showed that these patterns could be captured quantitatively.

Taken together, our results advance our understanding of human choice behavior in complex real-world environments. The results may also have broader implications for understanding consumer behavior. For example, we have found that customers frequently change to unhealthy food options after bad experiences. However, a potential strategy to increase the exploration of healthy food might be to increase healthy restaurants’ relative uncertainty by grouping healthy options with other frequently explored options such as Asian restaurants, which showed a comparatively lower relative uncertainty per restaurant.

While we have focused on using cognitive models to predict human choice behavior, the same issues come up for the design of recommendation engines in machine learning. These engines use sophisticated statistical techniques to make predictions about behavior, but do not typically try to pry open the human mind^28^. This is a missed opportunity; as models of human and machine learning have become increasingly intertwined, insights from cognitive science may help build more intelligent machines for predicting and aiding consumer choice.

## Acknowledgements

ES was supported by the Harvard Data Science Initiative. SJG was supported by the Office of Naval Research under Grant N000141712984.

## Author contributions statement

ES, RB, and BB extracted and analyzed the data. BCL, MTT and SJG supervised the work. All authors wrote the paper.

## Methods

### The Deliveroo data set

The data consisted of a representative random subset of customers ordering food from the online food delivery service “Deliveroo”. The data set contained 195,333 fully anonymized customers. These customers placed 1,613,968 orders over two month (February and March 2018) in 197 cities. There were 30,552 restaurants in total leading to an average of 155 restaurants per city. We arrived at this data set by filtering out customers with less than 5 orders (too little data points to analyze learning and exploration) and more than 100 orders (likely multiple people sharing an account).

### Clustering analysis

We removed the cuisine type “European” for this analysis as it was found to contain little information about customer choice behavior. This is unsurprising, given that manual cuisine type tags vary in quality and information content. Next, we analyzed for each cuisine type how much exploring this type on a time point *t* was predictive of exploring another cuisine type on a time point t + 1, using a linear regression model. Repeating this analysis for every combination of cuisine types lead to the graph shown in Figure 4a. We then analyzed the resulting matrix of *r*^2^-values by using hierarchical clustering.

### Similarity judgments

To elicit similarity ratings between different cuisine types, we asked 200 participants on Amazon’s Mechanical Turk to rate the similarities between two randomly sampled types out of the 20 types used for the clustering analysis reported above. Participants were paid $1 and had to rate 50 pairs of cuisine types in total. The study took less than 10 minutes on average.

## Supporting information

### Data set

We used the following variables to analyze customer’s exploration behavior: anonymized customer ID, anonymized restaurant ID, anonymized order ID, the name of the city an order was placed in, the cuisine type of the order (180 types in total), the standardized price of the restaurant indicating how much more expensive the restaurant was than a median restaurant within the same city (from 0.25 to 2), the standardized estimated delivery time of the order (z-scores from -2 to 3), how many orders a customer had placed previously, whether or not it was the first time a customer had ordered from the chosen restaurant, the mean rating of the restaurant at the time of the order from 4 to 5^1^, the number of previous ratings for the restaurant at the time of the order, and the eventual rating the customer provided (from 1 to 5). We also calculated every order’s RPE by subtracting the mean restaurant rating from the eventual order rating.

**Figure S1.**
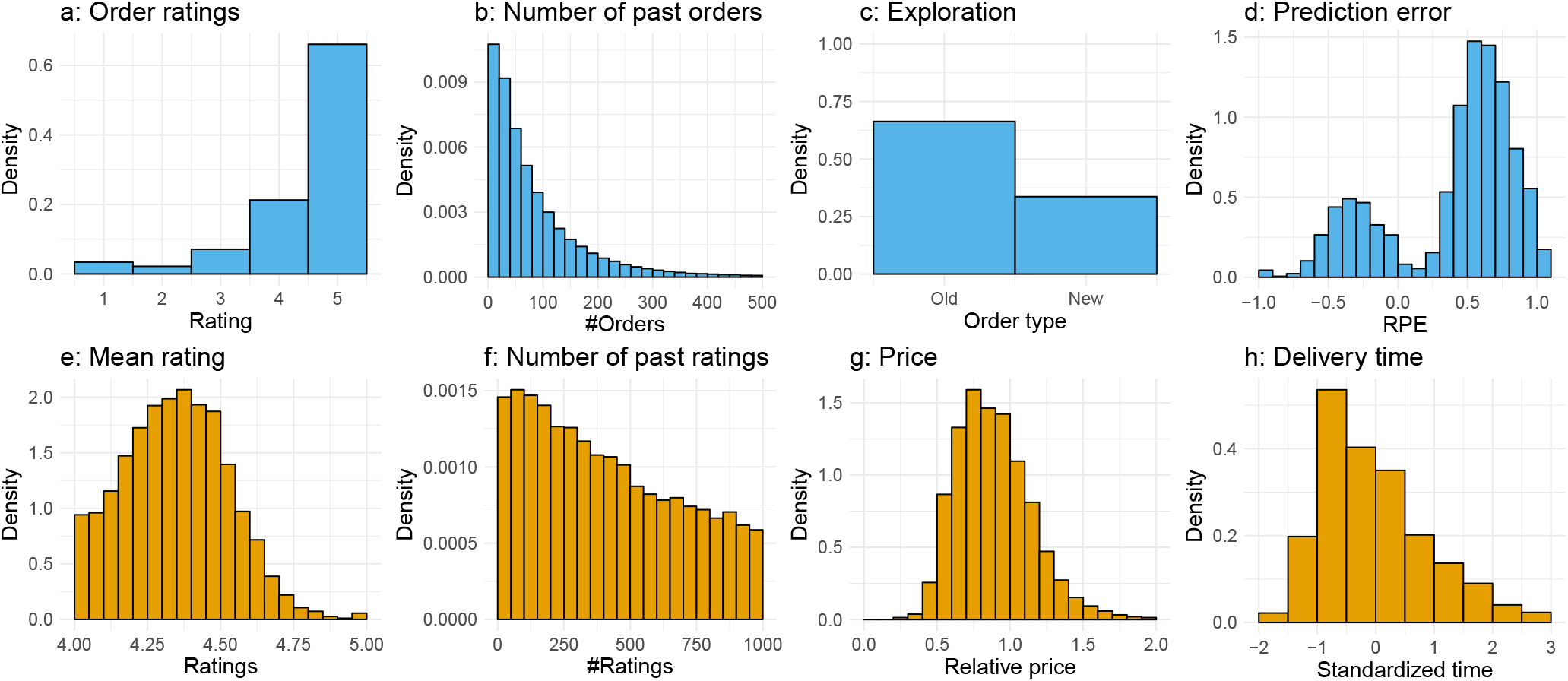
Distributions of customer-specific (blue) and restaurant-specific (orange) variables. **a:** Customers’ order ratings from 1 (low quality) to 5 (high quality). **b:** Number of past orders per customer. **c:** Proportion of choosing an old vs. a new restaurant. **d:** Prediction error, defined as the difference between the actual order rating and the mean restaurant rating. **e:** Mean restaurant ratings at the time of an order. Ratings have been truncated at a value of 4 to avoid instability induced by infrequent low average ratings. **f:** Number of past ratings per restaurant at the time of an order. **g:** Relative price per restaurant at the time of an order. **h:** Standardized delivery time at the time of an order.

The distributions of all variables are shown in Figure S1.

### Statistical tests

As our data set was large, almost any comparison would be significant at the *α* = 0.05-level. We therefore report the means and 99.9% confidence intervals for each group when reporting differences. We believe that this descriptive comparison makes the size of the differences more interpretable.

### Mixed-effects regression

We report the step-wise results for both mixed-effects regression analyses. We compare models based on their Akaike Information Criterion (AIC) and Bayesian Information Criterion (BIC).

#### Factors influencing exploration

For the first mixed-effects regression, we regressed a restaurant’s price (Price), mean rating (Rating), number of past ratings (#Ratings), and estimated delivery time (Time) onto whether or not a customer explored that restaurant. Additionally, we entered a random intercept for each customer.

The variables price, rating and the number of ratings all had a linear effect onto a restaurant’s probability of being explored, whereas the average delivery time had a nonlinear effect (the expression Time^2^ in Tab. S1 indicates that we entered both a linear and a quadratic effect of time into the final model). The final model contained all variables and had a fit of BIC=251772.

**Table S1.** Results of mixed-effects logistic regression analyzing determinants of exploration.

| Model | AIC | BIC | $\chi^2$ | $\text{Pr}( > \chi^2 )$ |
| --- | --- | --- | --- | --- |
| Intercept only | 258107 | 258127 | – | – |
| Price | 258064 | 258095 | 45 | <.001 |
| Price+Rating | 257294 | 257334 | 772 | <.001 |
| Price+Rating+#Ratings | 251784 | 251835 | 5511 | <.001 |
| Price+Rating+#Ratings+Time<sup>2</sup> | 251772 | 251843 | 16 | <.001 |

#### Signatures of directed exploration

For the second mixed-effects logistic regression, we regressed a restaurant’s value difference (Value), relative uncertainty (Uncertainty) and price (Price) onto the exploration variable. The value difference is defined as the difference in ratings for a given restaurant compared to the average of all restaurants within the same cuisine type. The relative variance is defined as the difference between a restaurant’s variance in ratings and the average variance per restaurant within the same cuisine type. The price is the relative price indicating how much more expensive a restaurant was compared to the city’s median restaurant price.

**Table S2.** Results of mixed-effects logistic regression analyzing signatures of directed exploration.

| Model | AIC | BIC | $\chi^2$ | $\text{Pr}( > \chi^2 )$ |
| --- | --- | --- | --- | --- |
| Intercept only | 258107 | 258127 | – | – |
| Value | 257184 | 257214 | 924 | <.001 |
| Value+Price | 257152 | 257193 | 34 | <.001 |
| Value+Price+Uncertainty | 257066 | 257117 | 88 | <.001 |

The final model contained all three variables and produced a fit of BIC=257117. Thus, relative uncertainty was a significant contributor to customers’ exploration behavior beyond value difference and relative price, a strong signature of directed exploration.

### Model comparison

#### New customers data set

For the model comparison, we created a data set containing only customers who had just started ordering food on the Deliveroo website (i.e., new customers). Moreover, we filtered out all customers who did not rate all of their orders. This resulted in a data set of 3,772 customers in total. We used this data set to compare different models of learning combined with different decision strategies, treating customers’ chosen restaurants as the arm of a bandit and their ratings as the resulting reward.

#### Gaussian Process

We use Gaussian Process (GP) regression as a Bayesian model of generalization. A GP is defined as a collection of points, any subset of which is multivariate Gaussian. Let 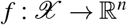 denote a function over input space 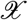 that maps to real-valued scalar outputs. This function can be modeled as a random draw from a GP:

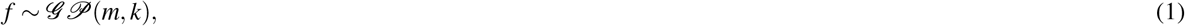

where *m* is a mean function specifying the expected output of the function given input **x**, and *k* is a kernel function specifying the covariance between outputs:

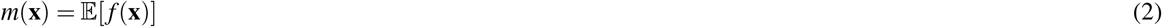

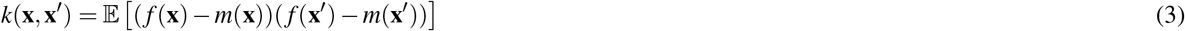

We fix the prior mean to the mean value of ratings within a given city and use the kernel function to model generalization over the restaurant-specific features.

Conditional on observed data 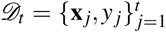, where 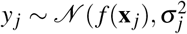 is drawn from the underlying function with added noise 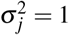, we can calculate the posterior predictive distribution for a new input **x**_*_ as a Gaussian:

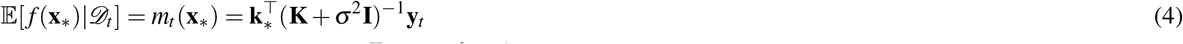

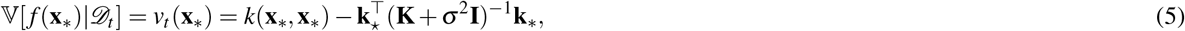

where **y** = [*y*_1_,…, *y_t_*]^⊤^, **K** is the *t* × *t* covariance matrix evaluated at each pair of observed inputs, and **k**_*_ = [*k*(**x**_1_, **x**_*_),…, *k*(**x**_*t*_, **x**_*_)] is the covariance between each observed input and the new input **x**_*_.

To model customers’ generalization over restaurants’ features, we assume that customers can use the presented features at the time of an order to predict a restaurant’s quality, i.e. how much they will like it. These features are the price, the mean rating, the number of past ratings, and the delivery time.

#### Radial Basis Function kernel

We use a Radial Basis Function (RBF) kernel as a component of the 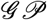 algorithm of generalization. The RBF kernel specifies the correlation between inputs **x** and **x′** as

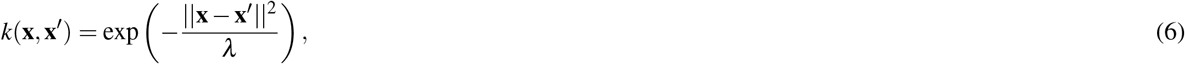

where *λ* is a length-scale parameter controlling function smoothness. This kernel defines a universal function learning engine based on the principles of Bayesian regression and can model any stationary function.

#### Bayesian Mean Tracker

The Bayesian Mean Tracker model is implemented as a Bayesian updating model, which assumes no temporal dynamics. In contrast to the GP regression model (which also assumes constant means over time), the Mean Tracker learns the rewards of each restaurant independently, by computing an independent posterior distribution for the mean *μ_j_* for each restaurant *j*. We implemented a version that assumes rewards are normally distributed (as in the GP model), with a known variance but unknown mean, where the prior distribution of the mean is a normal distribution. This implies that the posterior distribution for each mean is also a normal distribution:

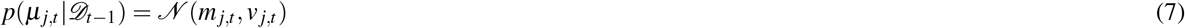

For a given option *j*, the posterior mean *m_j,t_* and variance *ν_j,t_* are only updated when it has been selected at trial *t*:

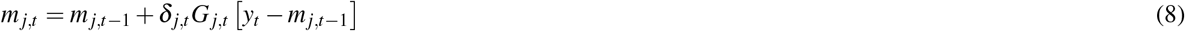

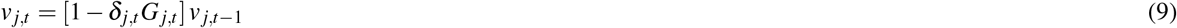

where *δ_j,t_* = 1 if option *j* is chosen on trial *t*, and 0 otherwise. Additionally, *y_t_* is the observed reward at trial *t*, and *G_j,t_* is defined as:

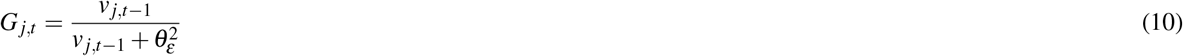

where 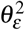 is the error variance, which we fixed to 0.2. Intuitively, the estimated mean of the chosen option *m_j,t_* is updated based on the difference between the observed value *y_t_* and the prior expected mean *m_j,t-1_*, multiplied by *G_j,t_*. At the same time, the estimated variance *ν_j,t_* is reduced by a factor of 1 − *G_j,t_*, which is in the range [0,1]. The error variance 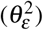 acts as an inverse sensitivity, where smaller values result in more substantial updates to the mean *m_j,t_*, and larger reductions of uncertainty *ν_j,t_*. We set the prior mean to the mean value of all restaurants within a city and the prior variance to *ν*_*j*,0_ = 5.

This model does not generalize at all and can therefore only learn about a restaurant’s quality by sampling it. Thus, it predicts that every novel restaurant will just be as good as the average of all restaurants in a city.

#### Sampling strategies

Given the normally distributed posteriors of the expected rewards, which have mean *μ* (**x**) and uncertainty (formalized here as standard deviation) *σ*(**x**), for each restaurant **x** (for the Mean Tracker, we let *μ*(**x**) = *m_j,t_* and 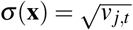, where *j* is the index of the restaurant characterized by **x**), we assess different sampling strategies that make probabilistic predictions about how much customers will like a given restaurant. In particular, we combine the Bayesian Mean Tracker with a mean-greedy sampling strategy and the Gaussian Process regression with both a mean-greedy and an upper confidence bound sampling strategy (details below).

#### Upper Confidence Bound sampling

Given the posterior predictive mean *μ*(**x**) and its attached standard deviation 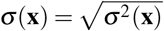, we calculate the upper confidence bound using a weighted sum

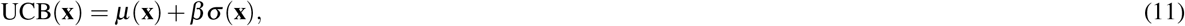

where the exploration factor *β* determines how much reduction of uncertainty is valued (relative to exploiting known high-value options). We fix *β* = 1 for our model comparison, indicating a tendency towards directed exploration.

#### Mean Greedy Exploitation

A special case of the Upper Confidence Bound sampling strategy (with *β* = 0) is a greedy exploitation component that only evaluates points based on their expected rewards

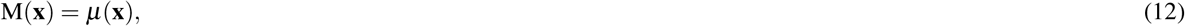

This sampling strategy only samples options with high expected rewards, i.e. greedily exploits the environment.

### Model comparison

We fit all models to a customers’ data until time point *t* and then make predictions about choices on time point *t* + 1. We apply a softmax choice rule to transform each model’s prediction into a probability distribution over options:

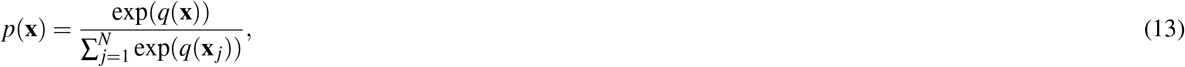

where *q*(**x**) is the predicted value of each option **x** for a given model (e.g., *q*(**x**) = UCB(**x**) for the UCB model).

#### One-step ahead prediction errors

We fit all models—per customer—to the data a customer has seen until time point *t* and then make forecasts about choices at time point *t* + 1. For example, the Gaussian Process model is fitted to all past restaurants a customer has sampled, using the restaurant’s features (i.e. prize, mean rating, number of ratings and delivery time) as the independent variables and the customer’s ratings as the dependent variable. Afterwards, it can be used to make predictions about other restaurants’ expected ratings (and uncertainties), that can be mapped onto probabilities. The difference between the Gaussian Process model and the Bayesian Mean Tracker is that the Bayesian Mean Tracker does not use any generalization over features, but only updates its predictions (which are equated to the overall mean at the beginning) by sampling a restaurant. The difference between the mean-greedy GP-M model and the GP-UCB model containing a directed exploration component is that the GP-M model equates a restaurant’s utility with the predicted mean rating, whereas the GP-UCB model equates a restaurant’s utility with its upper confidence bound.

Crucially, it is never possible to assess all restaurants a customer looked at and could have ordered from at a particular time point. We therefore compare the utility of the chosen restaurant to an average restaurant in the same city. For example, for the Gaussian Process model, we compared how much more likely a customer’s choice was compared to a restaurant with average feature values. For the BMT model, we compare the assessed utility to the overall average of restaurants in a city.

#### Predictive accuracy

The error of predictions (computed as predictive log loss) is summed up over all one-step ahead predictions, and is reported as *predictive accuracy*, using a pseudo-*R*^2^ measure that compares the total log loss for each model to that of a random model:

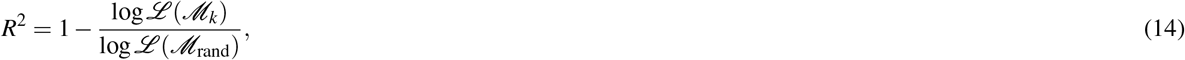

where 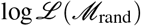 is the log loss of a random model (i.e., picking options with equal probability) and 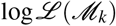 is the log loss of model *k*’s one-step-ahead prediction error. A *R*^2^ = 0 corresponds to a prediction accuracy equivalent to chance, while *R*^2^ = 1 corresponds to a theoretically perfect predictive accuracy, since 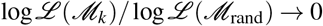 when 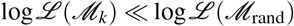.

### Sampling entropy and directed exploration

We calculated customers’ mean entropies over the next 4 samples after either a positive or a negative reward prediction error. Shannon’s entropy is defined as

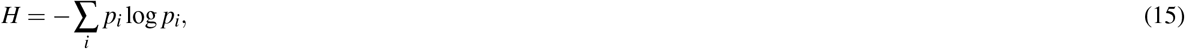

where *i* indicates a restaurant within customer’s 4 next choices. One of our predictions was that entropy would be higher after negative RPEs than after positive RPEs. We derived this prediction from the fact that a UCB sampling strategy updates both its mean and uncertainty after observing an outcome. After a bad experience, the mean and standard deviation both go down, whereas after a good experience the mean goes up but the standard deviation goes down. We confirmed this prediction in our data. Here, we check if this prediction holds in simulated data that was produced by either the GP-UCB or the GP-M model.

#### Synthetic data

For a first check of our prediction, we generated synthetic data using samples from a univariate Gaussian Process. Specifically, we created a one-dimensional meshed grid of options with *x* ∈ [0,0.2,0.4, ⋯, 10]. We then sampled a target function from a Gaussian Process with *λ* = 1 and optimized this function by using either a softmaximized mean-greedy (GP-M) or a upper confidence bound exploration strategy (GP-UCB) over 20 trials. Moreover, we tracked the models reward prediction error, defined as the difference between its predicted mean for a sampled option and the actual outcome of that option. We repeated this simulation 100 times for both models and afterwards calculated the sampling entropy for the models’ next 4 choices after having observed an outcome. Feeding the reward prediction error into a mixed effects regression with the sampling entropy as the dependent variable and a random intercept for simulation number, we found a significant effect of RPEs onto entropy for the GP-UCB model (*β* = −0.43, *SE* = 0.01, *t*(1454.5) = −32.70, *p* < .001), but not for the GP-M model (*β* = −0.001, *SE* = 0.006, *t*(1454.5) = −0.122, *p* = .9). Thus, UCB sampling leads to higher entropy after negative outcomes than after positive outcomes, whereas this difference is not pronounced in data generated by a soft-maximizing sampling strategy.

#### Customer data

In a second analysis, we looked at the sampling entropy difference between the GP-M and GP-UCB models in simulated customer choice data. We focused on the data for the new customers generated for our model comparison. For each customer, we generated a choice set by extracting all sampled restaurants and their features (price, rating, number of ratings, and delivery time). Furthermore, we estimated a utility distribution (the distribution of ratings for each restaurant by customer), by using a hierarchical model with a normal distribution, 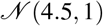, as the prior of the mean and a Cauchy distribution, Cauchy(0,1), as the prior over the variance of the restaurant’s utility. The resulting data set can be seen as 3,772 consecutive bandit tasks, where each task contains as many options as unique restaurants a customer had sampled and—for each restaurant—a reward distribution estimated by a hierarchical model based on that customer’s ratings. We then let both a GP-UCB and a GP-M model perform within this task, letting them sample as many restaurants as each of the customers had sampled. Afterwards, we calculated the entropy of the next 4 sampling steps as a function of the RPE (Fig. S2).

To assess the effect of RPE on sampling entropy, we regressed—for both sampling strategies individually—the RPE onto the entropy of the next 4 sampling steps in a mixed effects regression while also adding a random intercept for simulated customers. This showed that RPE had a significant effect onto sampling entropy for the UCB sampling strategy (*β* = 0.031, *p* = .007) but not for the softmax mean-greedy sampling strategy (*β* = 0.015, *p* = .07). We therefore conclude that our theoretical prediction holds in both synthetic and data-driven simulations of exploratory behavior over time.

### Clustering analysis

As described in the Materials and Methods, we clustered the 20 most frequent cuisine types appearing within our data set. One appropriate clustering solution of this analysis contained 7 main clusters. The scree plot of the clustering analysis (Fig. S3) confirmed our 7 clusters solution.

### Effect of environment analysis

To estimate whether or not customers explored more frequently in cities with higher mean ratings, we calculated the average rating over all restaurants as well as the average exploration rate for every city. The correlation between these two variables was *r* = 0.32, *t*(98) = 3.19, *p* = .002. Even when simultaneously correcting for a city’s average restaurant price, order volume, average number of ratings per restaurant, and average number of ratings per customer, the resulting partial correlation was still significant (*r* = 0.25, *t*(98) = 2.54, *p* = .02).

### Predictability analysis

For the predictability analysis, we assessed—for every city—how predictable customers’ ratings were based on the 4 features used throughout all of our analyses. Doing so, we only used the data set consisting of orders that customers had rated afterwards. We then sampled a learning and a test set for each city consisting of 100 orders each, fitted a linear regression model to the learning set, and used this model to predict customers’ order ratings in the test set. Repeating this analysis 100 times for every city revealed how predictable the quality of restaurants within one city was, measured by how well the regression model performed in the test set. We then correlated this predictability measure with the quality of exploratory choices (the mean rating of explored restaurants within a city). This correlation was significantly positive (*r* = 0.73, *t*(114) = 10.8, *p* < .001). Again simultaneously correcting for a city’s average restaurant price, order volume, average number of ratings per restaurant, and average number of ratings per customer, the partial correlation remained significant (*r* = 0.48, *t* (114) = 5.7, *p* < .001).

**Figure S2.**
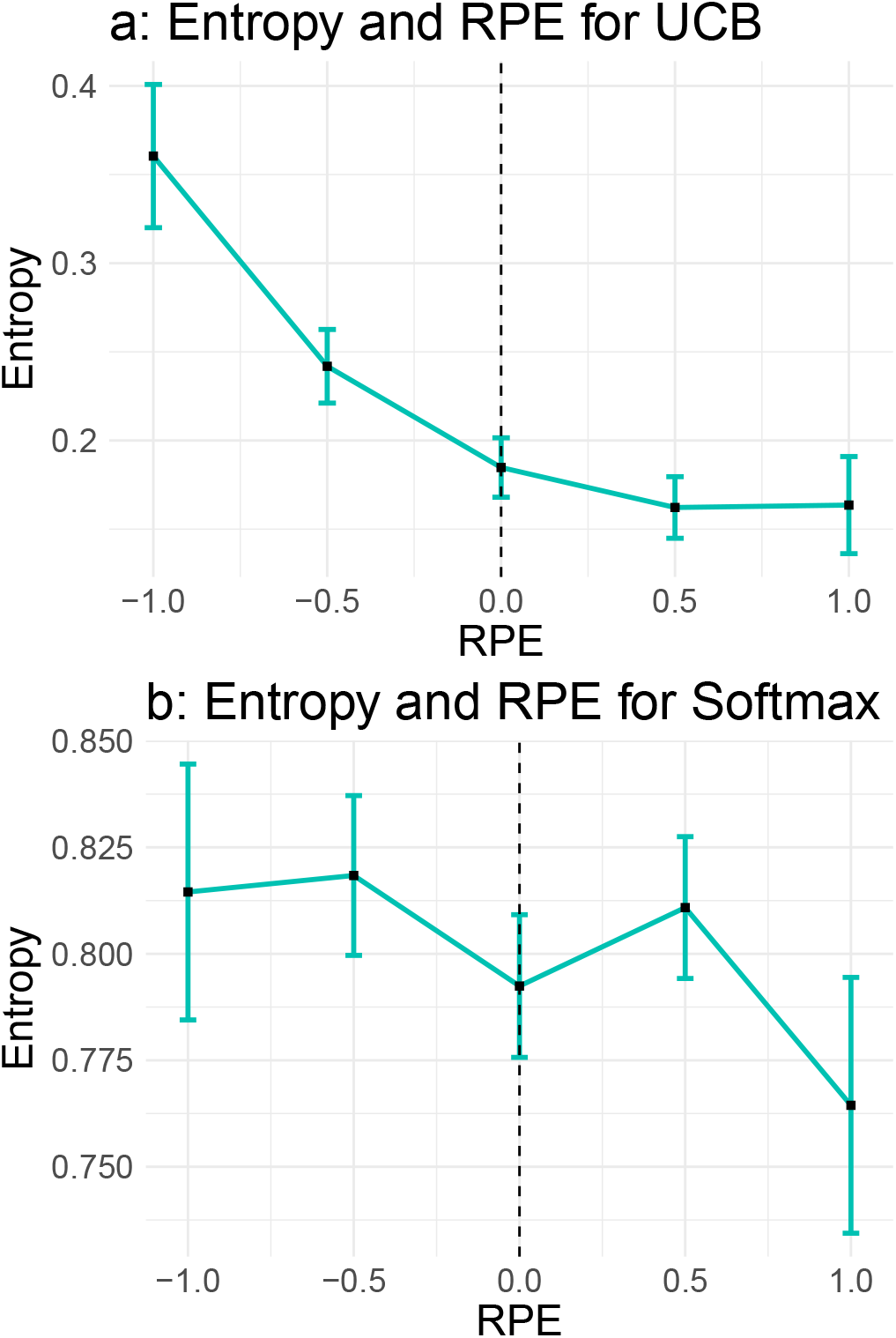
Entropy and prediction error for UCB and (softmaximized) mean-greedy sampling in a generated restaurant data set.

**Figure S3.**
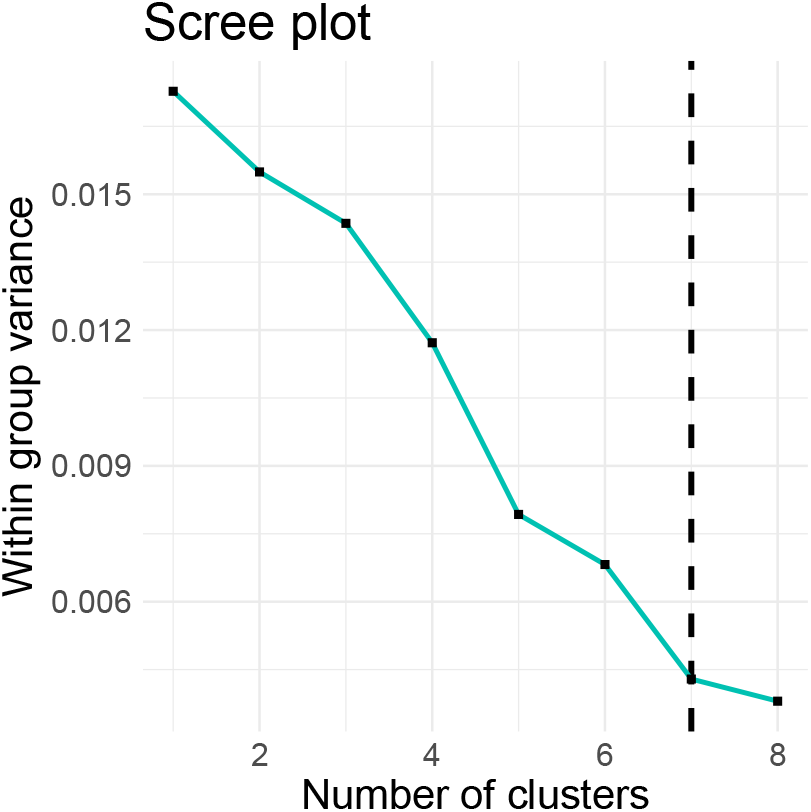
Scree plot of clustering analysis showing the amount of within variance by number of clusters. Dashed line indicate 7 clusters solution.

**Figure S4.**
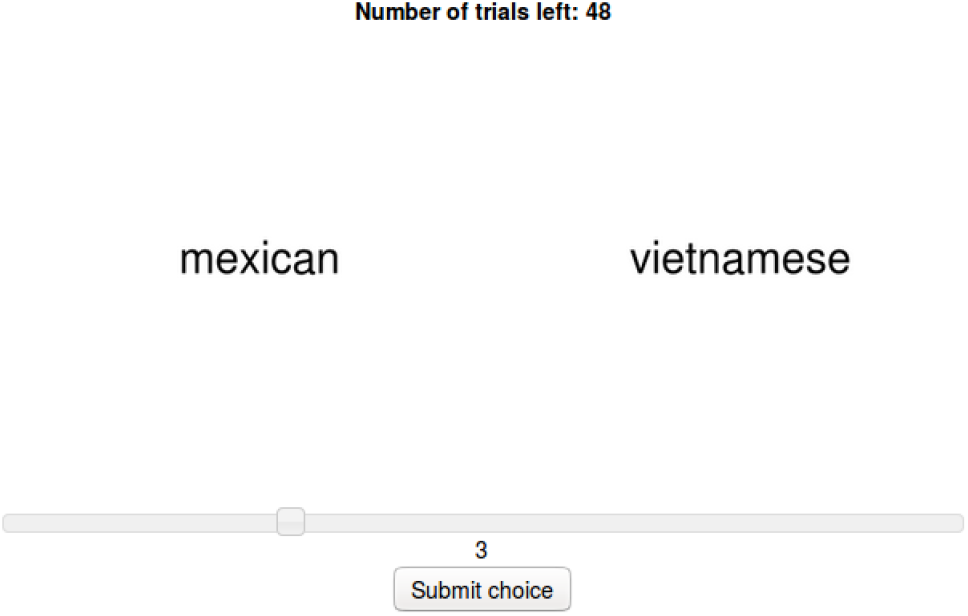
Screen shot of similarity rating study.

### Estimating causal directions

Three of our main theoretical predictions were: (1) customers explore more in cities with higher average ratings; (2) customers explore more successfully in cities where ratings were more predictable; and (3) relative uncertainty has a positive effect on customers’ tendency to explore a restaurant. Even though these predictions were derived from past empirical studies, it is nonetheless hard to make strong claims about causal directions in a natural and complex data set. Here, we additionally use another statistical method to estimate causal directions based on observational data proposed by Peters et al.^29^. Specifically, this method uses additive noise modeling to assess the residuals when performing a nonlinear regression from one variable to another and vice versa, and then applies kernel independence tests to decide about the causal direction, judging the direction as more likely in which the resulting residuals are more independent. We refer the interested reader to the original paper, but note here that this method gets up to 85% of classifications correct in a very challenging “causal directions benchmark” correct.

We applied this model to the city-specific variables of the mean exploration rate and the average restaurant rating. When regressing the exploration rate onto a city’s average restaurant rating using general additive models, this method assessed the probability of independence of the residuals as *p* = 0.46, whereas that probability was *p* = 0.58 the other way around. This method therefore weakly classified the average rating to be more likely the cause of the mean exploration rate than vice versa. We then used this approach to assess the directions of the connection between a city’s predictability and customer’s exploratory success. Whereas the probability of independence of the residuals was *p* = 0.08 when regressing exploratory success onto predictability, that probability was *p* = 0.85 the other way around. There was thus strong evidence that predictability caused exploratory success according to the causal direction estimation method.

Finally, we analyzed the effect between a restaurant’s relative uncertainty and the tendency to be explored. To do this, we estimated every customer’s mean relative restaurant uncertainty and mean proportion of exploratory choices. This analysis assessed if customers who explored more did so because of high relative uncertainty in their environment or if customers exploring more often caused higher uncertainties. The resulting p-values for independence were both relatively small as this was a very large data set for both regressions (10^-10^ and 10^-7^). However, the p-value for independence when regressing exploration onto relative uncertainty was 10^3^-times lower than vice versa, showing strong evidence that relative uncertainty led to increased exploration, according to the causal estimation model. Taken together, these results yielded additional evidence that the postulated directions of our predicted and confirmed effects are correct.

## Footnotes

1 In total, 94% of the restaurants had higher ratings than 4 and behavior for restaurants with an average rating lower than 4 was unstable. None of the main results change when analyzing the full data set, but estimates for this part of the space were unreliable.

## References

1. Whittle, P. Multi-armed bandits and the gittins index. J. Royal Stat. Soc. Ser. B (Methodological) 143–149 (1980).

2. Gershman, S. J. Deconstructing the human algorithms for exploration. Cognition 173, 34–42 (2018).

3. Speekenbrink, M. & Konstantinidis, E. Uncertainty and exploration in a restless bandit problem. Top. Cogn. Sci. 7, 351–367 (2015).

4. Frank, M. J., Doll, B. B., Oas-Terpstra, J. & Moreno, F. Prefrontal and striatal dopaminergic genes predict individual differences in exploration and exploitation. Nat. Neurosci. 12, 1062 (2009).

5. Auer, P. Using confidence bounds for exploitation-exploration trade-offs. J. Mach. Learn. Res. 3, 397–422 (2002).

6. Riefer, P. S., Prior, R., Blair, N., Pavey, G. & Love, B. C. Coherency-maximizing exploration in the supermarket. Nat. Hum. Behav. 1, 0017 (2017).

7. Wilson, R. C., Geana, A., White, J. M., Ludvig, E. A. & Cohen, J. D. Humans use directed and random exploration to solve the explore–exploit dilemma. J. Exp. Psychol. Gen. 143, 2074 (2014).

8. Cohen, J. D., McClure, S. M. & Angela, J. Y. Should I stay or should I go? how the human brain manages the trade-off between exploitation and exploration. Philos. Transactions Royal Soc. Lond. B: Biol. Sci. 362, 933–942 (2007).

9. Mehlhorn, K. et al. Unpacking the exploration–exploitation tradeoff: A synthesis of human and animal literatures. Decision 2, 191 (2015).

10. Kaelbling, L. P., Littman, M. L. & Moore, A. W. Reinforcement learning: A survey. J. Artif. Intell. Res. 4, 237–285 (1996).

11. Thrun, S. B. Efficient exploration in reinforcement learning. (1992).

12. Sutton, R. S., Barto, A. G., Bach, F. et al. Reinforcement learning: An introduction (MIT press, 1998).

13. Srinivas, N., Krause, A., Kakade, S. M. & Seeger, M. W. Information-theoretic regret bounds for gaussian process optimization in the bandit setting. IEEE Transactions on Inf. Theory 58, 3250–3265 (2012).

14. Daw, N. D., O’Doherty, J. P., Dayan, P., Seymour, B. & Dolan, R. J. Cortical substrates for exploratory decisions in humans. Nature 441, 876 (2006).

15. Findling, C., Skvortsova, V., Dromnelle, R., Palminteri, S. & Wyart, V. Computational noise in reward-guided learning drives behavioral variability in volatile environments. bioRxiv DOI:10.1101/439885 (2018).

16. Dezza, I. C., Angela, J. Y., Cleeremans, A. & Alexander, W. Learning the value of information and reward over time when solving exploration-exploitation problems. Sci. reports 7, 16919 (2017).

17. Knox, W. B., Otto, A. R., Stone, P. & Love, B. The nature of belief-directed exploratory choice in human decision-making. Front. Psychol. 2, 398 (2012).

18. Gershman, S. J. Uncertainty and exploration. Decision (in press).

19. Gershman, S. J. & Tzovaras, B. G. Dopaminergic genes are associated with both directed and random exploration. Neuropsychologia 120, 97–104 (2018).

20. Wu, C. M., Schulz, E., Speekenbrink, M., Nelson, J. D. & Meder, B. Generalization guides human exploration in vast decision spaces. Naure Hum. Behav. 2, 915–924 (2018).

21. Schulz, E., Franklin, N. T. & Gershman, S. J. Finding structure in multi-armed bandits. bioRxiv 432534 (2018).

22. Badre, D., Kayser, A. S. & D’Esposito, M. Frontal cortex and the discovery of abstract action rules. Neuron 66, 315–326 (2010).

23. Wimmer, G. E., Daw, N. D. & Shohamy, D. Generalization of value in reinforcement learning by humans. Eur. J. Neurosci. 35, 1092–1104 (2012).

24. Gershman, S. J. & Niv, Y. Novelty and inductive generalization in human reinforcement learning. Top. Cogn. Sci. 7, 391–415 (2015).

25. Schulz, E., Konstantinidis, E. & Speekenbrink, M. Putting bandits into context: How function learning supports decision making. J. Exp. Psychol. Learn. Mem. Cogn. 44, 927–943 (2018).

26. Stojic, H., Schulz, E., Analytis, P. P. & Speekenbrink, M. It’s new, but is it good? how generalization and uncertainty guide the exploration of novel options. (2018).

27. Schultz, W., Dayan, P. & Montague, P. R. A neural substrate of prediction and reward. Science 275, 1593–1599 (1997).

28. Griffiths, T. L. Manifesto for a new (computational) cognitive revolution. Cognition 135, 21–23 (2015).

29. Peters, J., Mooij, J. M., Janzing, D. & Schölkopf, B. Causal discovery with continuous additive noise models. The J. Mach. Learn. Res. 15, 2009–2053 (2014).

